# Offline coil position denoising enhances detection of TMS effects

**DOI:** 10.1101/256081

**Authors:** Leonardo Claudino, Sara J Hussain, Ethan R Buch, Leonardo G Cohen

## Abstract

**OBJECTIVE:** Transcranial magnetic stimulation (TMS) is extensively used in basic and clinical neuroscience. Previous work has shown substantial residual variability in TMS effects even despite use of on-line visual feedback monitoring of coil position. Here, we aimed to evaluate if off-line denoising of variability induced by neuronavigated coil position and orientation deviations can enhance detection of TMS effects.

**METHODS:** Retrospective modeling was used to denoise the impact of common neuronavigated coil position and rotation deviations during TMS experimental sessions on motor evoked potentials (MEP) to single pulse TMS.

**RESULTS:** Neuronavigated coil deviations explained approximately 44% of total MEP amplitude variability. Offline denoising led to a 136.71% improvement in the signal to noise ratio (SNR) of corticospinal excitability measurements. CONCLUSIONS: Offline modeling enhanced detection of TMS effects by removing variability introduced by neuronavigated coil deviations.

**SIGNIFICANCE:** This approach could allow more accurate determination of TMS effects in cognitive and interventional neuroscience.

**HIGHLIGHTS:**

- Coil deviations impact TMS effects despite use of on-line neuronavigation feedback.
- Offline denoising of coil deviation impacts on TMS effects significantly reduced variability at trial level.
- Offline denoising also significantly improved overall SNR of TMS effects.

## 1. INTRODUCTION

Transcranial magnetic stimulation (TMS) is widely used in cognitive and interventional neuroscience (Dayan et al., 2013). TMS effects are variable within and between individuals (Herrmann et al., 2006, Wassermann, 2008, Pasley et al., 2009, Nicolo et al., 2015), as shown by the stochastic peak-to-peak amplitudes of motor evoked potentials (MEPs) to single TMS pulses (Kiers et al., 1993, Zarkowski et al., 2006, Goldsworthy et al., 2016). Biological sources of variability in TMS effects include the composition of underlying brain and non-brain tissues exposed to the stimulation field (Chen et al., 2010, Opitz et al., 2011, De Geeter et al., 2012, Bestmann, 2015), the distribution of corticospinal output pathways recruited by an individual TMS stimulus (Di Lazzaro et al., 2001, Di Lazzaro et al., 2012, Di Lazzaro et al., 2014) and habituation of neuron populations (Brighina et al., 2009, Pitkanen et al., 2017).

One important non-biological factor that contributes to variability in TMS effects is coil position and orientation (Mills et al., 1992, Di Lazzaro et al., 2001). Frameless stereotactic neuronavigation systems partially addressed this problem by providing realtime information about the position and orientation of the coil relative to the individual subject’s scalp- or brain-defined target (Ruohonen et al., 2010). This information has been used to provide online visual feedback to experimenters, thus assisting in manual correction of TMS targeting errors due to coil or head movement at the time stimulation is delivered (Lioumis et al., 2009, Cincotta et al., 2010, Jung et al., 2010, Richter et al., 2013). Integration of robotic coil holders with neuronavigation systems can further improve targeting precision by using this feedback to adjust coil position and orientation in real-time (Lancaster et al., 2004). However, adoption of these robotic systems across TMS research laboratories has been minimal, most likely due to their high cost (Ambrosini et al., 2018). Overall, variability in TMS effects due to coil positioning remains a substantial problem in spite of online neuronavigation (Cincotta et al., 2010, Richter et al., 2013). It would be important to develop strategies to reduce noise introduced by trial-by-trial differences in coil position on TMS effects.

Here, we propose a novel offline denoising approach to address this problem. It uses trial-by-trial coil position deviations, commonly available in current neuronavigation systems but unused in subsequent statistical analyses, to retrospectively model and remove their impact on TMS effects.

## 2. METHODS

### 2.1. Participants

20 healthy adults participated in this study (13M, 6F; age=30±7.1yrs, range=22–48yrs). Neuronavigation data from three subjects was corrupted, resulting in 18 usable datasets. Subjects provided written informed consent and the study was approved by the NIH Combined Neuroscience IRB. Healthy status was verified prior to study participation via neurological examination and brain MRI performed by trained clinicians.

### 2.2. TMS and MEP recording

Subjects were seated in an armchair during the study. Monophasic TMS was delivered with a hand-held figure-of-eight coil (oriented approximately 45° relative to the mid-sagittal line) attached to a Magstim 200^2^ unit (Magstim, Inc). Electromyography (EMG) was recorded at 5kHz using Signal software (CED, Cambridge UK) from the left first dorsal interosseous (FDI) using disposable adhesive electrodes arranged in a belly-tendon montage.

#### 2.2.1. Hotspot estimation

The left FDI hotspot was estimated as the scalp position most reliably eliciting the largest MEPs following suprathreshold stimulation. The stimulation intensity was initially set to 50% maximum stimulator output and increased if this intensity was insufficient for generating an observable muscle twitch. A grid-like search was carried out centered around location C4 of the International 10-20 System. The search was carried out by initially moving the coil in approximately 2cm increments over the grid, which covered an approximate area spanning 4cm anterior and lateral to 4cm posterior and medial of location C4. Between 2-3 TMS stimuli were delivered on average at each location. Hotspot candidates identified during the initial search were then refined further by testing adjacent locations 1cm anterior, posterior, medial and lateral to each candidate (6±4; median ± IQR). The median number of total TMS stimuli delivered during this procedure per subject was 118 ± 54.5 (median ± IQR).

#### 2.2.2. Resting Motor Threshold (RMT) estimation

The resting motor threshold (RMT) was estimated using an adaptive threshold-hunting algorithm (Awiszus, 2003) based on parameter estimation by sequential testing (PEST; MTAT 2.0; http://www.clinicalresearcher.org/software.htm).

#### 2.2.3. MEP recordings

A total of 600 suprathreshold TMS stimuli (120% RMT; interstimulus interval=5 ± 0.75s) were then delivered to the FDI hotspot, except for one subject to whom only 557 TMS stimuli were delivered due to time constraints. A total of 10,757 MEP trials from 18 subjects were recorded. None of these trials were excluded from analyses.

### 2.3. Neuronavigation

Frameless stereotactic neuronavigation (Brainsight 2, Rogue Research) was used to localize subject head, FDI hotspot target, and TMS coil position and orientation within the same spatial reference frame. Real-time visual feedback of the TMS coil position and orientation relative to the FDI hotspot target was used to guide hand-held TMS coil positioning over the course of the experimental session. Coil position and orientation deviations (i.e. – spatial and rotational disparity between coil and FDI hotspot target location, Fig.1) were acquired at the time of TMS delivery so 10,757 coil deviation entries (one for each recorded MEP) were also recorded.

**Figure 1.**
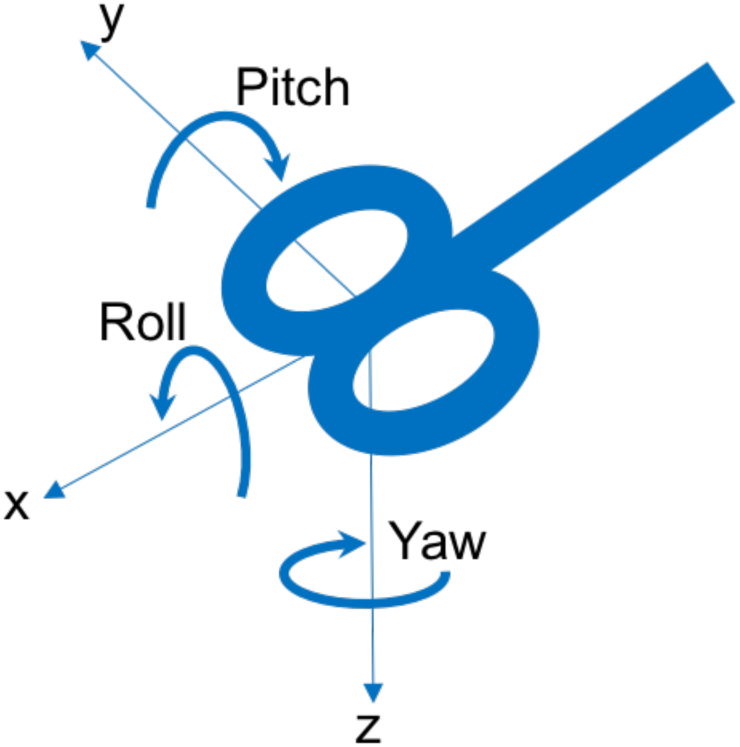
Coil position and orientation deviations. For each TMS pulse, the neuronavigation software recorded the displacement of the coil with respect to the estimated hot spot scalp location as a homogeneous transformation matrix (rotations and translations). Later, as part of the offline analysis, these matrices are condensed into spatial (x, y and z) and rotational (roll, pitch, yaw) coordinates, as depicted.

### 2.4. Data analysis

#### 2.4.1. Pre-processing

For each trial, peak-to-peak motor evoked potentials (MEP) amplitudes were calculated as the difference between the maximum and minimum voltage deflections between 20–40ms post-stimulus. Each 6-D coil deviation coordinate (x, y, z, yaw, pitch and roll; Fig.1) was normalized by its interquartile range (IQR) across all trials. Next, we used a greedy heuristic to increase the density of MEP amplitudes per coil deviation coordinate value (see Supplemental materials for details). Finally, to increase the density of MEP amplitudes per coil deviation coordinate value, we independently re-quantized each coordinate, one by one, into a number of bins (within 1 to 1000) that maximized detection of significant changes in MEP amplitudes with respect to the bin that included coordinate 0 (no deviation; Sign test, p<0.05, Bonferroni-corrected). See Supplemental materials for details.

#### 2.4.2. Denoising

The relationship between the coil deviation (relative to the FDI hotspot target) and MEP amplitude was modeled on a trial-by-trial basis using a mixed effects model with random intercepts at the subject level (Model 1):

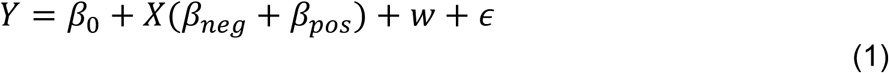

where *Y _T_*_×1_ is the matrix with all *T* trials, *X _T_*_×6_ is the design matrix of fixed effects with each row consisting of all 6-D coil deviations for each trial. We used a piecewise linear model centered at 0 to account for directional contributions of coil deviations to MEP amplitudes, represented as the fixed-effect slopes, *β_neg_* and *β_pos_*. Vector *w* is the individual subject’s mean MEP with respect to the grand mean MEP, *β*_0._ Residuals *∊* are assumed to be Gaussian-distributed with zero-mean and unknown variance.

Since *w* reflects *any* between-subjects difference and not just those differences explained by coil deviations, a second piecewise multiple linear regression was used to obtain the fraction of between-subjects differences accounted for by *only* coil deviations (Model 2):

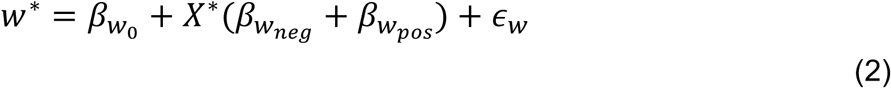

where *w\** expresses the between-subjects difference as a linear function of coil deviation. *X\** represents the individual subject median coil deviations reduced to *K* dimensions. Dimensionality reduction is necessary because a full coil deviation model at the subject level would rely on too many parameters (piecewise 6-D × 2 + intercept = 13 parameters) given current sample size (N=18). Here, dimensionality reduction is achieved via principal component analysis (PCA) based upon singular value decomposition (SVD):

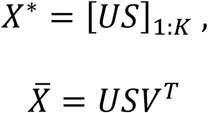

where 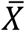 is a matrix with each subject’s median 6-D coil deviations minus the median MEP amplitude between subjects. *X\** is obtained by (1) projecting 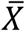 onto the linear subspace spanned by singular vectors selected after running a bi-directional elimination step regression procedure (R’s *step* function with default arguments, keeping the model with the lowest Akaike Information Criterion (AIC) score) and (2) augmenting the number of rows from N to T by replicating each subject’s entry by the number of trials recorded for that subject. Residuals *∊_w_* are assumed to be normal with zero-mean and unknown variance. Slopes *β*_[*w_neg,_w_pos_*]_ represent the contribution (relative to the intercept 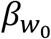) of individual subject’s median coil deviation to the between-subjects MEP difference *w*\* at negative and positive values of these coordinates,.

Models 1 and 2 are combined by substituting *w* in Eq.1 with *w\** (Eq.2) yielding a single expression of raw MEP amplitudes:

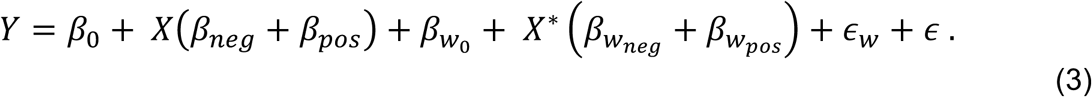

Finally, denoised MEP amplitudes, *Ŷ*, were obtained by subtracting the total fraction of MEP variability explained by coil deviations from raw MEP amplitudes (Y, Eq. 3) yielding:

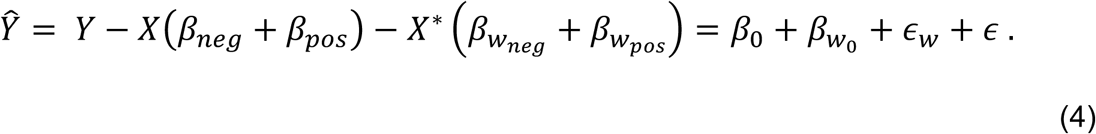

Note from the above equation that the denoising procedure can result in an upward or downward adjustment to measured individual trial MEP amplitudes. The direction of this adjustment is dependent upon: (1) the coil deviation (X) observed on that particular trial, and (2) the individual subject’s median coil deviation relative to the median deviation between subjects (X*) weighed by the fixed and random effects β parameters as determined from model fits.

#### 2.4.3. Assessment of denoising

We assess the effect of denoising as percent changes in median, interquartile range (IQR) and signal-to-noise ratio (SNR) from raw (*Y*) to denoised (*Ŷ*) MEP amplitudes:

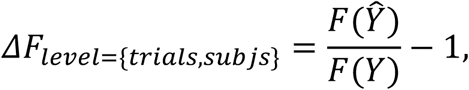

where *F* is one of the following functions: *Median, IQR* or *SNR* and the level of analysis is either trials (*level* = *trials*) or subjects (*level* = *subjs*).

## 3. Results

Neuronavigated coil deviations varied across trials and subjects (Fig. 2), and explained approximately 44% of total MEP amplitude variance (Conditional R^2^ = 0.4445; (Nakagawa et al., 2013).

**Figure 2.**
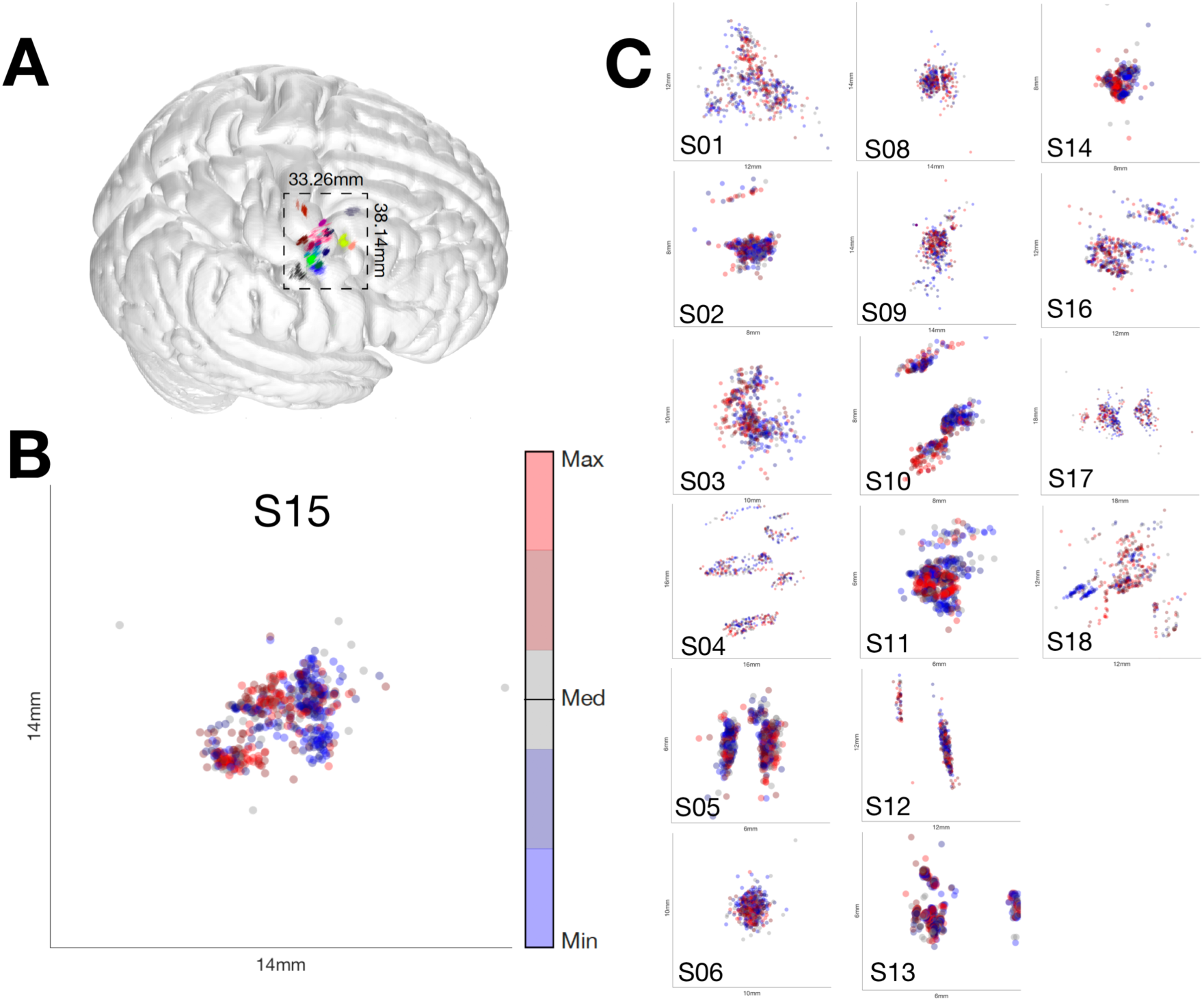
Neuronavigated coil deviations contributed to MEP amplitude variability. (A) TMS coil peak stimulation field location was projected onto the cortical surface using 6D coil positions. Nonlinear spatial registration (FSL FNIRT) was used to map individual subject data onto the MNI152 brain to visualize differences in estimated hotspot locations and within-subject neuronavigated TMS coil deviations. Dots depict the estimated stimulation field peak location on the cortical surface for all individual trials and subjects (different colors) in MNI space. Estimated hotspot locations varied across subjects. Percentile ranked MEP amplitude as a function of cortical surface stimulation field peak location for one enlarged representative (B) and all other (C) subjects. Blue (0-40 percentile MEP amplitude) and red (60-100 percentile MEP amplitude) clusters in the maps indicate the contribution of neuronavigated coil deviations to MEP amplitude variability. Please note that one subject (S07) was omitted because T1 MRI was not acquired.

Removal of MEP amplitude variability caused by the neuronavigated coil deviations using this denoising method resulted in improvements in SNR at both trial-by-trial and subject levels. Fig. 3A depicts trial-by-trial raw and denoised MEP amplitude distributions. Trial-by-trial SNR improved by ΔSNR_trials_=136.71 *%* (one-tailed random permutation test, p = .00005) explained by a decrease in variability of ΔIQR_trials_=26.23% (one-tailed random permutation test, p = .00005) and an increase in median MEP of ΔMedian_trials_=74.62% (one-tailed random permutation test, p = .00005, Fig 3B). All permutation tests used 20,000 repetitions.

**Figure 3.**
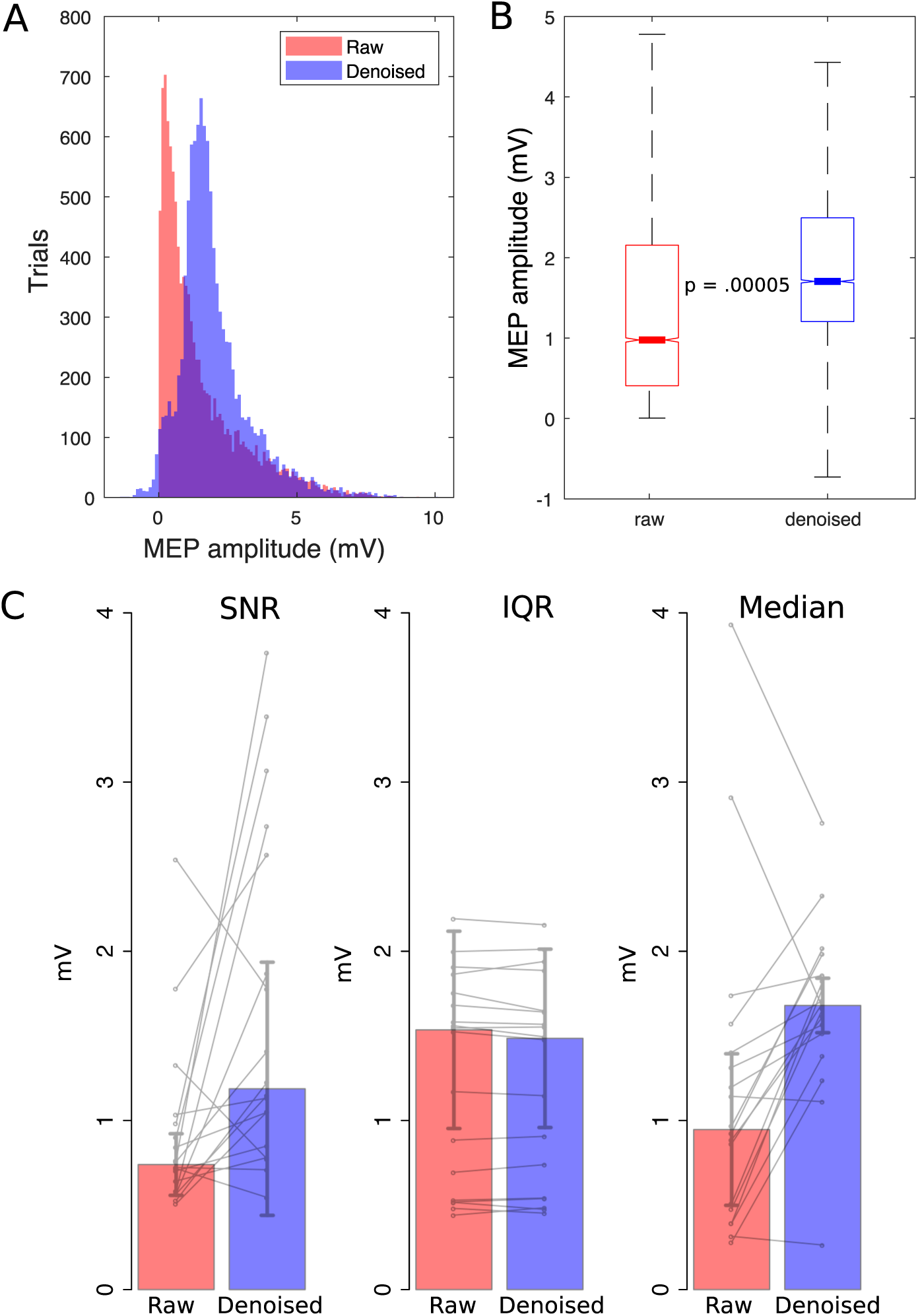
Signal-to-noise ratio across (A) and within (B) subjects. **A.** Trial-by-trial raw and denoised MEP amplitude distributions. Denoising resulted in a more symmetric distribution of MEP amplitudes. **B.** Trial-by-trial median and interquartile range (IQR, box) of raw and denoised MEP amplitudes. Denoising resulted in higher median and smaller IQR leading to ΔSNR_trials_=136.71% improvement in signal-to-noise ratio (SNR). **C**. Raw and denoised SNR, IQR and median MEP amplitudes at the subject level. Note the ΔSNR_subjs_=65.39 % improvement in SNR accounted largely by increases in median MEP amplitudes in the absence of differences in IQR. Increases in SNR were observed in 14 of the 18 subjects tested (solid lines).

Between subject SNR improved by ΔSNR_subjs_=65.39% (Wilcoxon sign rank test, V=150, p = .001682) explained predominantly by an increase in median MEP of ΔMedian_subjs_=65.29% (Wilcoxon sign rank test, V=152, p = .001167, Fig 3C). Between subject variability did not differ (ΔIQR_subjs_= .9986%, Wilcoxon sign rank test, V=76, p = .3509). The SNR improvement was evident in 14 out of the 17 subjects tested.

## 4. Discussion

The main finding of this study was that retrospective modeling of coil position and orientation deviations improved signal-to-noise ratio (SNR) of TMS effects measured as the peak-to-peak amplitude of motor evoked potentials (MEP). It is possible that the observed improvements in SNR resulting from denoising could translate to better characterization of TMS effects. A similar approach has been taken in the field of neuroimaging (Nielsen et al., 2018) to address head motion – a source of noise analogous to coil deviation – which is known to contaminate BOLD signal modeling (Caballero-Gaudes et al. 2017), secondary estimates of functional (Power et al., 2012) and structural (Baum et al., 2018) connectivity, and the relationship between these indirect measures of brain activity with behavior (Siegel et al., 2016). Here, we found that neuronavigated coil deviations explained approximately 44% of total MEP amplitude variance, which is similar in magnitude to reported effects of head motion on BOLD fMRI resting-state functional connectivity measurements (Nielsen et al., 2018). The application of our proposed denoising procedure resulted in an increase in SNR by approximately 136.71%, which again is similar to SNR increases observed following denoising of head motion effects in fMRI data (Nielsen et al., 2018).

Trial-by-trial coil position and orientation deviation information saved after customary online neuronavigation was used to characterize the partial contribution of these factors to MEP amplitude variability, and then denoise MEP measurements offline. The proposed technique was designed to remove contributions of coil deviation and is intended to work with commonly available laboratory neuronavigation systems, where data required to carry out this analysis are typically available, but commonly unused in subsequent statistical analyses. While denoising clearly improved signal-to-noise ratios, it should be kept in mind that variance related to any other factor that strongly correlates with coil deviation could be removed by linear regression-based denoising as well.

Mechanical constraints can be used to stabilize coil (i.e. – mechanical arms or coil holders) and head position (i.e. – specialized chair head rests, chin rests or bite bars) and reduce relative deviations over the course of an experiment. However, such equipment may not be available in all TMS laboratories, or may not be compatible with experimental designs that include behavioral tasks. Integrating robotic coil holders that use neuronavigation system feedback to adjust coil position and orientation in real-time can improve accuracy (Lancaster et al., 2004), but is expensive and not commonly available (Ambrosini et al., 2018). In the end, none of these approaches will fully guarantee that the relative relationship between coil and head position and orientation remains stable over the duration of a given experiment. Our offline denoising approach could be used in combination with any of these mechanical approaches to address residual coil deviations that may occur.

Finally, it would be interesting to test this approach on previously acquired datasets where trial-by-trial neuronavigated TMS coil position data is available. Such data sets could include neurophysiological (i.e., motor threshold, intracortical inhibition (Hallett, 2007, Lioumis et al., 2009)) or behavioral (i.e., effects of TMS on reaction time (Chen et al., 1998, Johansen-Berg et al., 2002)) outcomes as dependent variables. In such datasets, βs (i.e. - β_[neg,pos,wneg,wpos]_ from Eqs. 1 and 2) can be obtained over all trials and subjects given the measured coil deviation from the estimated hotspot. The trial-by-trial denoised Ŷs for these datasets can then be calculated from these βs using Eq. 4. In the future, it will be interesting to study if denoising could reduce variability of TMS effects characterized across interventions, operators or locations (Julkunen et al., 2009, Fleming et al., 2012).

## 5. Conclusions

Offline neuronavigated coil deviation denoising may be a simple and feasible approach to enhance detection of TMS effects in cognitive and interventional neuroscience.

## Acknowledgments

This work was supported by the Intramural Research Program of the National Institute of Neurological Disorders and Stroke.

## Supplemental materials

### Binning of coil deviation coordinates

<u>The</u> binning pre-processing procedure used (Section 2.4.1) is a simple greedy heuristic similar to typical dimension-by-dimension feature selection techniques in machine learning (Hall, 2000) except that here the goal is to reduce sparsity. We depart from the premise that our coil deviation data is naturally oversampled (high resolution) so we really want to undersample. But we also want to avoid disruptive effects of heavy undersampling, which we do by monitoring the detection of MEP changes as a function of the number of bins using significance tests (Fig S1). Please note that these tests play no role in choosing trials or subjects in the analyses of median, IQR or SNR of MEPs and as such they do not compromise any reported statistics (see Fig. S2 for a comparison with results of when denoising was done without binning).

**Figure S1.**
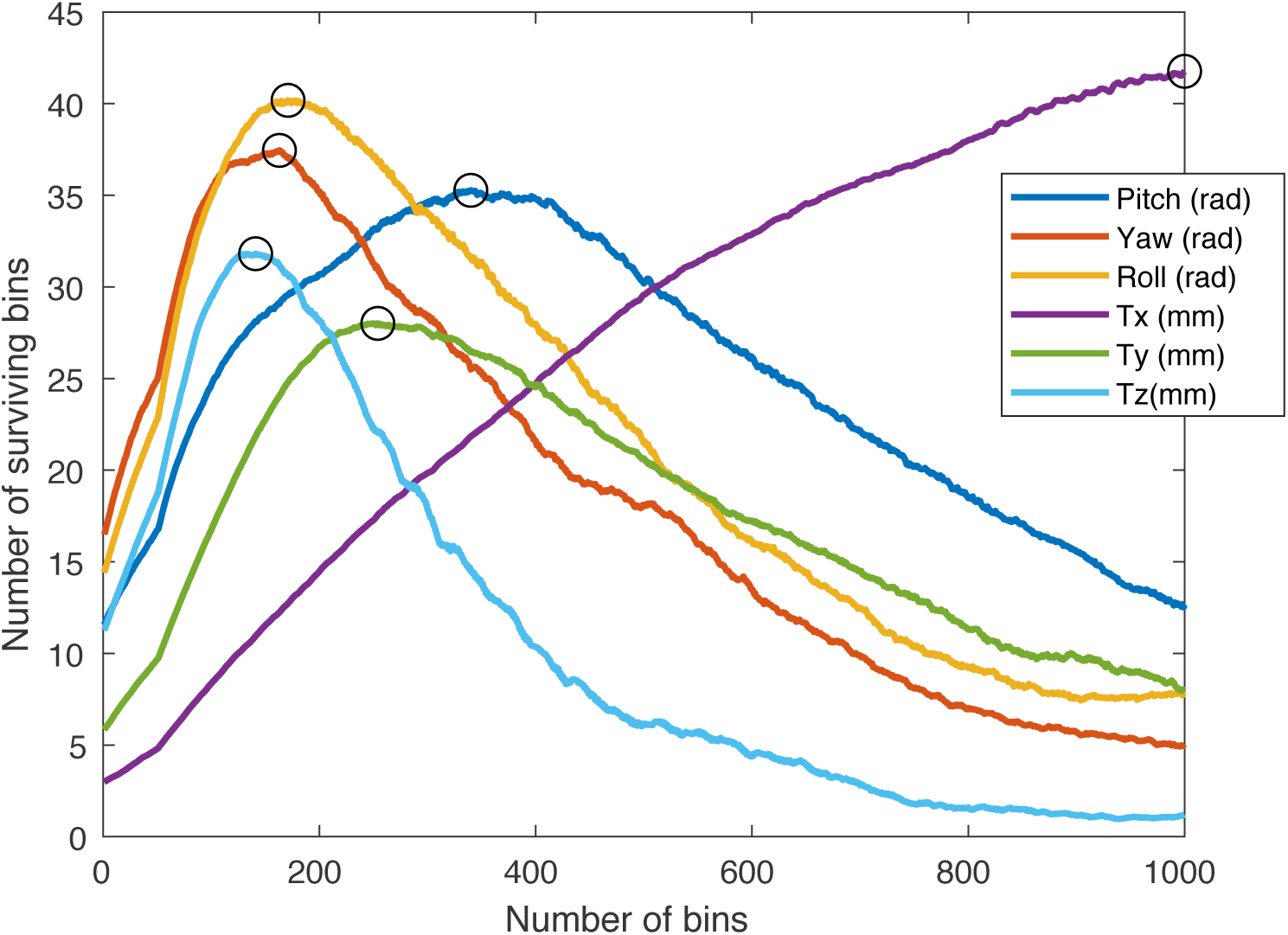
Greedy selection of number of bins for all coil deviation dimensions (different colors). For every candidate number of bins from 1 to 1000 (x axis) we calculate the subset of that number that produces significant changes in MEP from coil position = 0 (surviving bins, y axis). As we move along the x-axis, this number grows to a maximum (optimal) value (except for Tx, purple, that did not converge but was close enough to maximum). Choosing values lower (too much undersampling) or higher (too much oversampling) than the estimated optimal could hinder the ability to detect changes in MEP from coil deviation that would need to be accounted for. Note that different coil deviation dimensions can have different optimal number of bins (circles at each maxima). The curve was smoothed with a moving average filter of window length=100 to facilitate detection of maxima (movmean function in Matlab).

**Figure S2.**
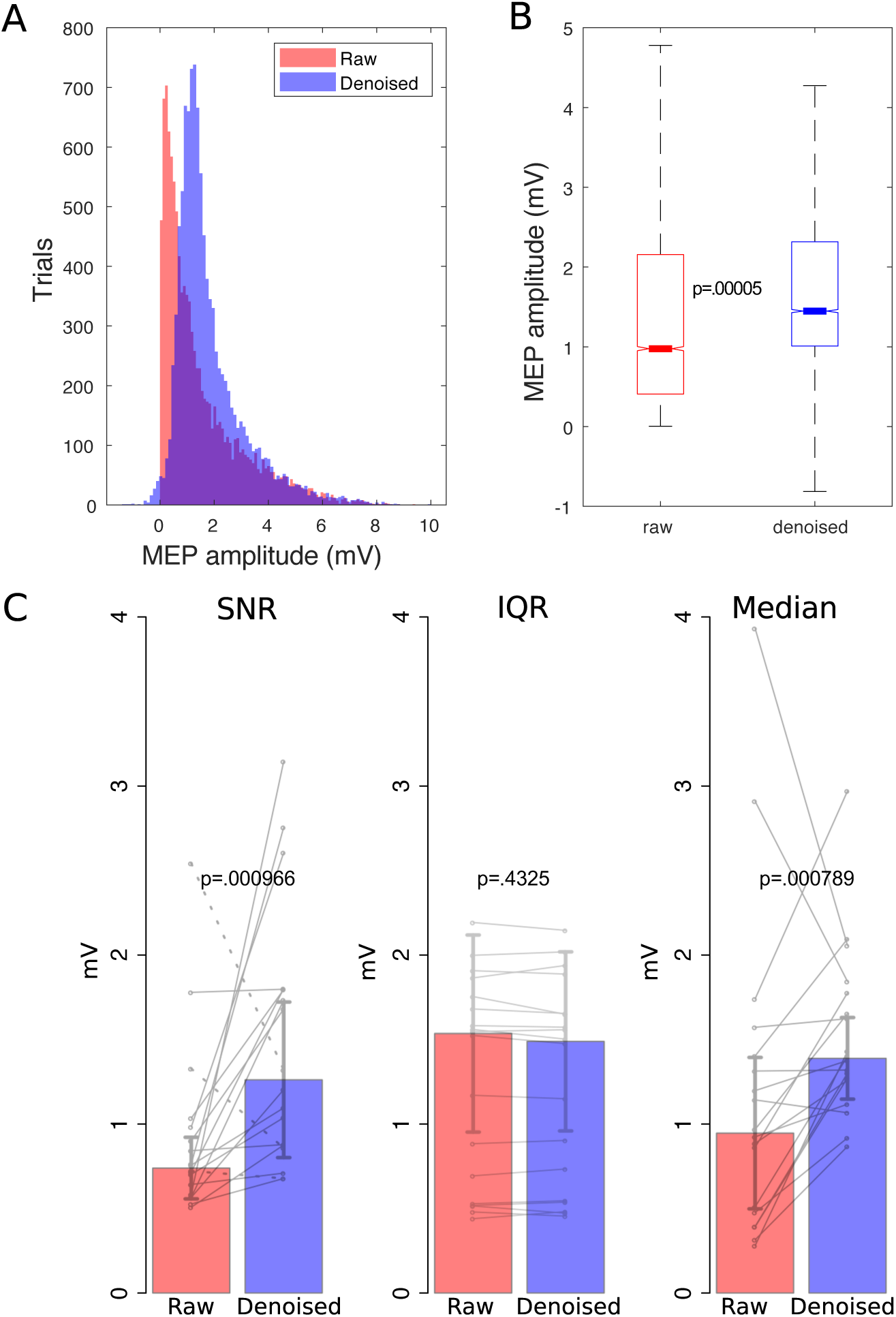
Denoising without binning. (A) Trial-by-trial raw and denoised MEP amplitude distributions. Results of denoising at trial level (B) and at the level of subjects (C) without binning. The shapes of denoised MEP distributions with and without binning are visually very similar (compare A above with Fig.3A). Not using binning produced similar changes in median (ΔMedian_trials_=48.26% and ΔMedian_subjs_=60.09%) IQR (ΔIQR_trials_=25.28% and ΔIQR_subjs_=.0.82%) and lower, and still substantially improved SNR (ΔSNR_trials_=98.42% and ΔSNR_subjs_=60.26%) when compared with when binning was used (Figs. 3B and 3C and Section 3).

